# Swimming Prevents Memory Impairment by Increasing the Antioxidant Defense in an Animal Model of Duchenne Muscular Dystrophy

**DOI:** 10.1101/751461

**Authors:** Priscila Mantovani Nocetti Ribeiro, Adriano Alberti, Viviane Freiberger, Letícia Ventura, Leoberto Ricardo Grigollo, Cristina Salar Andreau, Rudy José Nodari Junior, Daniel Fernandes Martins, Clarissa M. Comim

## Abstract

Duchenne muscular dystrophy (DMD) is a genetic disease which is associated to a progressive skeletical muscle degeneration. Swimming is usually indicated for avoiding impact and facilitating adherence because of a better adaptation to a warm water invironment and also for its benefits on cognition, and modulating memory and learning processes and for increasing antioxidant defenses in oxidative stress. The objective of this study was to evaluate the effects of a swimming protocol on memory and oxidative stress in an animal model of Duchenne muscular dystrophy. Methods: male mdx and wild type mice within 28 days were used in this study. The animals were trained in an stepped swimming protocol for four consecutive weeks. Twenty four hours after the last exercise day, aversive memory and habituation memory tests were performed and removed the encephalic structures of striatus, pre frontal cortex, hippocampus, and cortex and gastrocnemius and diafragma muscles to evaluate protein carbonilation and lipid peroxidation and free thiols. Results: it was verified that swimming was able to reduce significantly the levels of lipid peroxidation and protein carbonilation in gastrocnemius and hippocampus and striatus in exercised animals. Swimming has also prevented lipid peroxidation in diafragma. Besides, this swimming protocol was able to increase free thiols in gastrocnemius, diafragma and in analysed SNC structures. These results showed that swimming prevented aversive and habituation memory in mdx mice.

## Introduction

Duchenne muscular dystrophy (DMD) is one of the recessive X-linked PMS and mainly affects males. DMD usually presents in early childhood, characterized by delays in motor milestones. DMD leads to weakening of the skeletal muscles and is known to be the most common and severe type, due to its early onset and the rapid evolving of symptoms [1–3].

Treatment of patients diagnosed with Duchenne muscular dystrophy (DMD) needs to be multidisciplinary, careful, and always focused on the well-being of the patient. Given this, the role of physical therapy becomes crucial to the success of treatment, as it has achieved good results in the short term, such as maintenance of the autonomy of these individuals [4,5]. Another method that has been used to treat DMD is physical exercise [6,7]. Studies have been using physical exercise to decrease muscle deterioration, muscle contractures, bone fractures, and increase the time of functional independence, in individuals with DMD [8,9].

Swimming is one of the aerobic exercise modalities that has become very popular. Aerobic exercise in water is a viable alternative to exercise on land, because water, through its physical properties such as thrust, hydrostatic pressure, and viscosity, among others, brings increased well-being and quality of life [10]. Swimming, like walking and running, has health benefits compared to a sedentary lifestyle [11]. Swimming is generally indicated because with heated water it avoids impact and facilitates adherence by better adaptation to the environment [11]. Swimming is further described in the literature as having beneficial effects on patients: on cognition by modulating memory and learning processes; and in oxidative stress, by increasing antioxidant defenses [11]. Thus, the present study aims to evaluate the effects of swimming on memory and oxidative stress in an animal model of DMD.

## Materials and methods

### Animals

Male mdx and wild-type (WT) mice of the C57BL / 6 strain, 28 days old, weighing between 18 and 23 g from USP - São Paulo, were used. The animals were kept in the LANEX Vivarium and per box five animals were packaged, in a 12-h light/dark cycle (06:00 to 18:00), with food and water ad libitum. The environment was maintained at a temperature of 22 ± 2 °C. The study was carried out at LANEX and the Laboratory of Biochemistry and Molecular Biology during 2016 and 2017. This project was approved by the Animal Use Ethics Committee - CEUA of UNISUL under the number 16.003.4.01.IV.

### Experimental draw

The animals were divided into four groups of eight animals each: (1) unexercized WT; (2) exercised WT; (3) mdx not exercised; and (4) mdx exercised. Groups 2 and 4 were submitted to the swimming type aerobic exercise protocol for four weeks. Twenty four hours after the last day of training, aversive memory and habituation tests were performed. Afterward, the animals were sedated and submitted to assisted painless death procedure and the following structures were removed: gastrocnemius; quadriceps; diaphragm; prefrontal cortex; cerebellum; hippocampus; striatum; and cerebral cortex, for determination of lipid peroxidation, protein carbonylation, and thiol grouping.

### Swimming type aerobic exercise protocol

The groups of exercised animals were submitted to an aerobic swimming exercise protocol in a plastic container adapted for this purpose (170 x 100 mm), with 35 liters of water at 28 to 30 °C, divided into eight lanes. There were four consecutive weeks of exercise, four times a week, with daily sessions of 15 minutes in the first week, 20 minutes in the second week, and 30 minutes in the third and fourth week [12]. Of baby shampoo, 1 ml was used throughout the container to decrease surface tension of water [13]. After the protocol, the animals were gently dried. The groups of non-exercised animals did not perform any type of physical exercise, remaining in their housing boxes during the entire study period.

Exercise intensity was determined in the fourth week of protocol in WT and mdx animals. Blood samples were collected before the test and at the 10^th^ and 30^th^ minutes of exercise for subsequent analysis of lactate concentration. The criterion for considering intensity was the increase in concentration of no more than 1 mmol / L between the 10^th^ and 30^th^ minute of physical exercise. Hygienization of the blood collection site was performed with alcohol (70%). After this procedure, the distal portion of the tail of the animal was slightly sectioned with surgical scissors and 25 μl and a drop of blood was inserted in the lactate collection tape, and then, by means of a portable lactimeter, the blood level was measured. Before each test, the equipment was calibrated according to the manufacturer’s instructions [14].

### Aversive Memory Test

This test consisted of an acrylic box in which the floor was formed by parallel metal bars. A platform 7 cm wide and 2.5 cm long was placed near the left wall of the appliance. In the training session, the animals were placed on the platform and the time in seconds was measured for the animal to descend with all four legs from the platform. This time is called latency. Immediately after descending from the platform, the animal received a 0.2 mA shock for 2 s. In the test session, the animal was again placed on the platform and the time it took to descend (latency) the platform was measured, but no shock was given. The test was also terminated if the animal did not descend within a maximum of three minutes [15].

### Habituation memory test

This test was performed in a 40 x 60 cm open field, with four 50 cm high walls delimiting the area; three of them wood and one of transparent glass. The open field floor was divided into 16 equal squares, marked by black lines. In the training session, the animals were carefully placed in the square of the back left corner of the apparatus, from which they freely explored the environment for five minutes. Immediately afterward, the animals were returned to the box. The test session was held 24 hours after training, and the training procedure was then repeated. The number of four-legged crossings (crossings: motor activity) across the black lines and the number of times animals reared on their hind legs (rearings: exploratory activity) were evaluated in both sessions [16].

### Oxidative stress measures

Measurement of thiobarbituric acid reactive substances (TBARS) This method is used to evaluate the oxidation state of hydroperoxides in biological systems. Membrane lipid damage is determined by the formation of lipoperoxidation byproducts (such as MDA or malondialdehyde), which are substances reactive to thiobarbituric acid heating that are formed during peroxidation in membrane systems. MDA reacts with thiobarbituric acid (TBA), generating a pinkish-colored product read in a 535 nm microplate reader. The technique consists of the following form: firstly, the dilution value was calculated so that the TBARS reaction tube has 100 μl of tissue protein in 500 μl of BHT buffer. Thereafter, 500 μl of the 0.67% TBA solution was added. The tubes were placed in a dry bath at 96 °C for 30 minutes. To stop the reaction, the samples were placed on ice for 5 minutes. Finally, 200 μl of the reaction mixture was placed in 96-well microplates and read in the 535 nm microplate reader.

### Measurement of oxidative damage in proteins

This method was used for protein oxidation dosing. It is based on the principle that several ROS attack protein residues, such as amino acids, produce products with the carbonyl group, which can be measured by reaction with 2,4-dinitrophenylhydrazine. The carbonyl content is determined by a 370 nm microplate reader as described in the study by Levine et al. (1990). Firstly, the tissue was homogenized in 1 ml BHT buffer. Samples were centrifuged for 15 min at 4 °C at 14,000 rpm. Of the sample, 200 μl was separated to blank and 200 μl was separated for the test. Of 20% trichloro acetic acid (TCA), 100 μl was placed in all eppendorfs. It was centrifuged for 5 min at 14,000 rpm. The supernatant was discarded. The pellet was redissolved in 100 μl of 0.2 molar NaOH. Of dinitrophenylhydrazine 2.4 (DNTP), 100 μl was placed in the sample and allowed to stand for 1 h. Of 20% tTCA, 100 μl was placed in all eppendorfs, which were centrifuged for 3 min. The supernatant was then discarded. The pellet was washed 3 times with 500 μl ethanol and ethyl acetate (1: 1). For each wash, it was centrifuged for 3 min at 14,000 rpm and the supernatant discarded. After discarding the last wash, 1 ml of 3% sodium hydroxide (NaOH) was placed in all eppendorfs. The samples were taken to the bath at 60 °C for 30 minutes and read on the microplate reader at 370 nm.

### Thiol groupings

Sulfhydryl radicals represent all groups of thiols found in proteins such as albumin and in low molecular weight compounds such as glutathione. These groups may be oxidized when the oxidative stress is elevated. Determination of total sulfhydryl groups, protein-bound sulfhydryl groups, and sulfhydryl groups in low molecular weight compounds (free sulfhydryls) can be carried out using Ellman’s reagent (2,2-dinitro-5,5- dithiobenzoic acid – DTNB). The thiol groups react with DTNB to form a light-absorbing complex at 412 nm. The technique consists of adding 10% TCA to the same sample volume (1: 1 dilution). Prepared blank containing 100 μl TCA was added to 100 μl PBS. It was centrifuged for 15 minutes at 3,000 rpm (temperature 4 °C); the supernatant was collected and at 75 μl of this, 30 μl DTNB (1.7 mM) and 300 μl hydrochloric acid (TRIS-HCl) were added. It was allowed to react for 30 minutes and transferred to a 96-well plate. The samples were read in a 412 nm microplate reader.

### Protein Dosages

Proteins were determined by the BCA method and bovine serum albumin was used as standard. The method was based on the reaction of copper with proteins in basic medium. Samples were analyzed by a 562 nm plate reader.

### Statistical analysis

Data were entered into an electronic database in the IBM SPSS Statistics 24.0 software (@copyright IBM corporations and its licensors 1989, 2016). The Shapiro-Wilk normality test was applied to verify the behavior of the data. The data related to the open field habituation test were expressed as mean and standard deviation because they are parametric data. For differences between groups, a two-way analysis of variance (ANOVA) with Bonferroni post-hoc test was used. For differences between training and test in the same group, we used Student’s *t*-test for paired samples. The inhibitory avoidance test data were expressed as median and interquartile range because they are nonparametric data. Wilcoxon test was used for analysis between training and test in the same group. Biochemical test data were expressed as mean and standard deviation because they are parametric data. The two-way ANOVA with Bonferroni post-hoc test was used for analysis between groups. Data were considered statistically significant when *p* <0.05.

## Results

### Lactate measure

Figure 3 shows the results obtained after the measurement of lactate in the blood taken from the animals during the swimming protocol.

**Figure 3.**
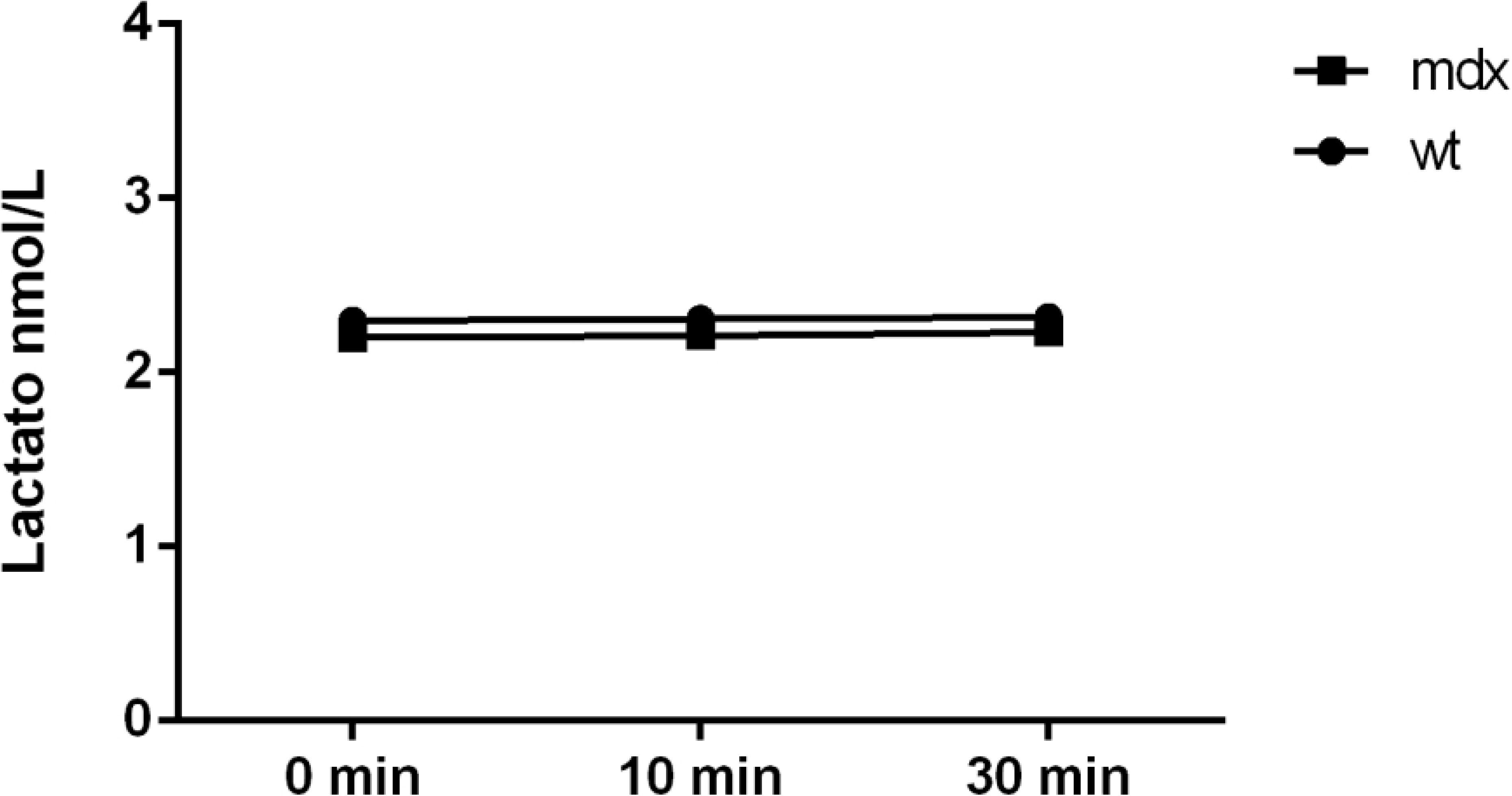
Lactate measurement of mdx and wt animals submitted to the swimming protocol.

Figure 3 shows that in both mdx and WT animals submitted to the swimming protocol, there was no change between the 10^th^ and 30^th^ minutes of exercise in blood lactate above 1 mmol / L in relation to rest values, being mean lactate values of 2.31 nmol / l for WT animals and 2.21 for mdx animals (i.e., remained below the lactate threshold).

Thus, the swimming protocol used in this study can be considered to be of moderate intensity.

### Learning and memory evaluation

Figure 4 shows the results obtained after the application of the swimming protocol on aversive memory (Fig. 4A) and habituation (Fig. 4B). Figure 4A shows the results of habituation memory evaluation through the open field test. It can be observed that there was no difference in the number of crossings and rearings (*p* > 0.05) between groups during the training phase, demonstrating that there was no difference in locomotor activity between the groups. The non-exercised and exercised WT animals showed significant changes between training and testing, both in the number of crossings and the number of rearings (*p* < 0.05), that is, there was no impairment of the evaluated memory. The group of non-exercised mdx animals (DMD) did not show significant changes between training and tests in the number of crossings and rearings (*p* > 0.95), evidencing a memory impairment. In contrast, the group of mdx animals that underwent the protocol showed a decrease in the number of crossings and rearings between training and testing (*p* < 0.05), suggesting a possible prevention of aversive memory impairment in mdx mice.

**Figure 4.**
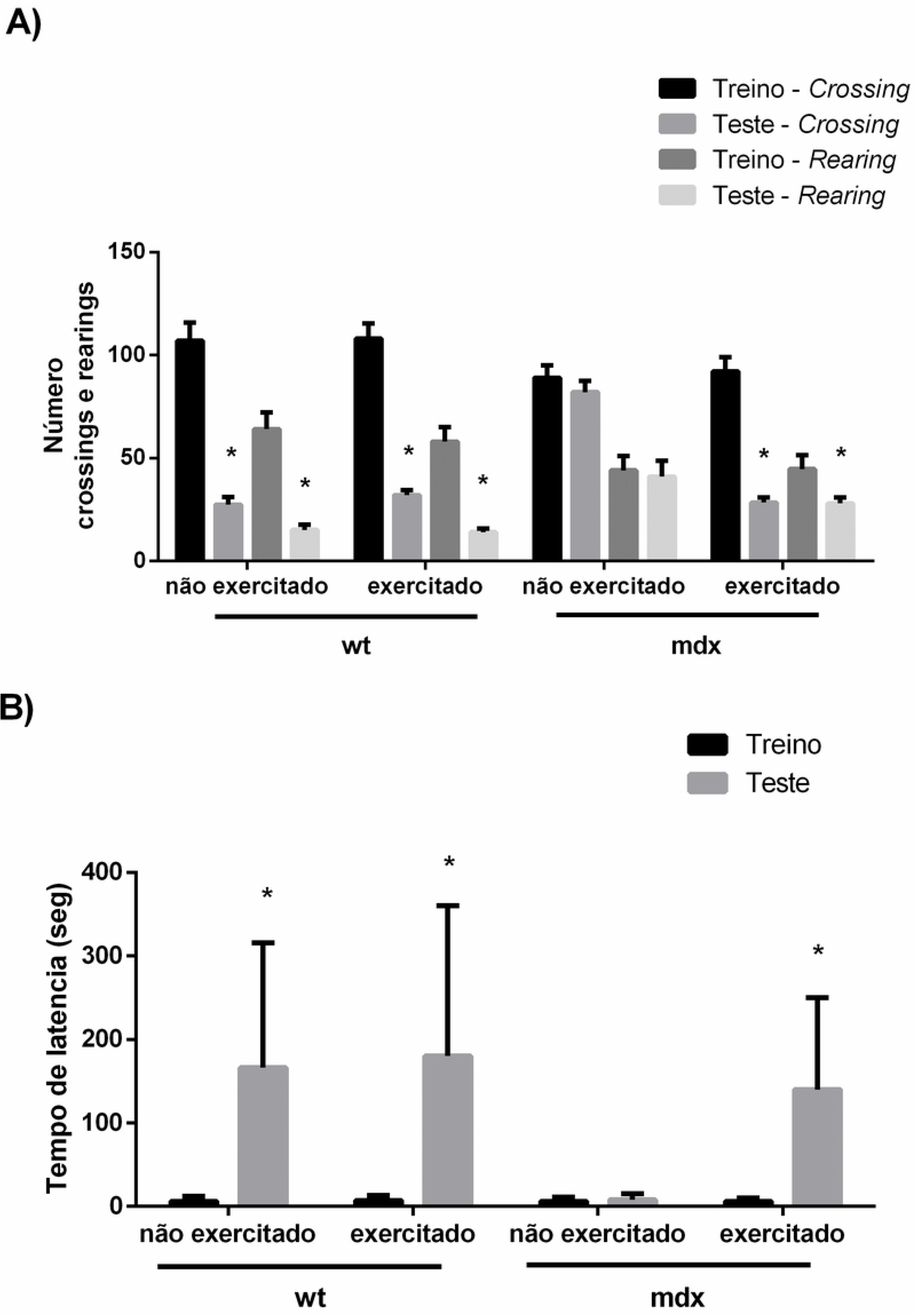
Effect of a swimming protocol on aversive and habituation memory of mice.

The results of aversive memory assessment by the inhibitory avoidance test are shown in Fig. 4B. In the group of non-exercised and exercised WT animals, there was a statistically significant difference in the latency time between training and testing, showing no impairment in aversive memory (*p* < 0.05). In the group of non-exercised mdx animals, there was no statistically significant difference between training and testing, evidencing an impairment of aversive memory (*p* > 0.05). The mdx animals submitted to the experimental protocol showed a statistically significant difference between training and testing, i.e., there was no impairment of aversive memory in these animals (*p* < 0.05).

### Evaluation of oxidative stress in the gastrocnemium muscle

Figure 5 shows the results of using a swimming protocol on lipid peroxidation (Figure 5A), protein carbonylation (Figure 5B), and free thiols (Figure 5C) in the gastrocnemius muscle.

**Figure 5.**
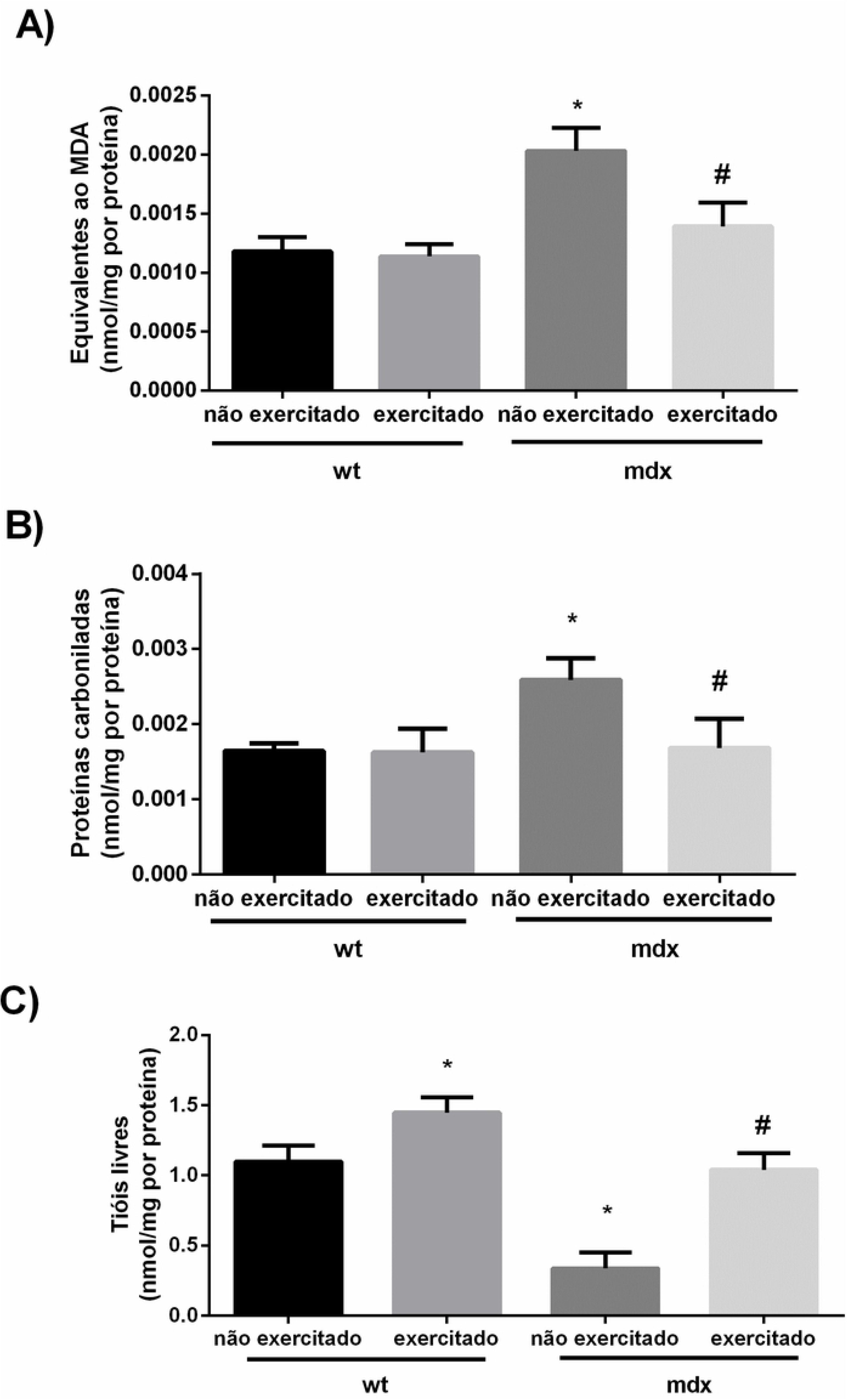
Effect of a swimming protocol on oxidative stress in mouse gastrocnemius muscle. (* when compared to the non-exercised wt group; ** when compared to the nonexercised mdx group).

Figure 5A shows the result of the assessment of lipid peroxidation in gastrocnemius. Unexercised mdx animals showed significantly higher levels of lipid peroxidation in gastrocnemius when compared to untrained wild animals (*p* <0.05). The mdx animals that underwent the experimental protocol presented a significant reduction of these levels when compared to the non-exercised mdx animals (*p* <0.05), showing that the experimental protocol used in this study protected against the increase of lipid peroxidation in gastrocnemius in mdx animals. It was observed that the non-exercised mdx animals showed a significant increase in protein carbonylation in gastrocnemius when compared to the non-exercised wild animals group (*p* <0.05). After the experimental protocol, the mdx animals showed significantly lower protein carbonylation levels when compared to the non-exercised mdx animals (*p* <0.05), demonstrating that the four-week swimming protocol was able to prevent the carbonylation increase. of protein observed in the gastrocnemius muscle of mdx animals (Figure 5B). The quantification of free thiols in gastrocnemius is shown in Figure 5C. It can be observed that there was a significant increase in the amount of free thiols in gastrocnemius of the group of wild animals submitted to the experimental protocol, when compared to the non-exercised wild animals (*p* <0.05). There was also a significant decrease in non-exercised mdx animals as compared to non-exercised wild animals (*p* <0.05). When the mdx animals underwent the experimental protocol, there was an increase in the amount of free thiols when compared to the unexercised mdx animals.

### Assessment of oxidative stress in diaphragm muscle

Figure 6 shows the results of a training protocol on lipid peroxidation (Figure 6A), protein carbonylation (Figure 6B), and several pounds (Figure 6C) in the diaphragm muscle.

**Figure 6.**
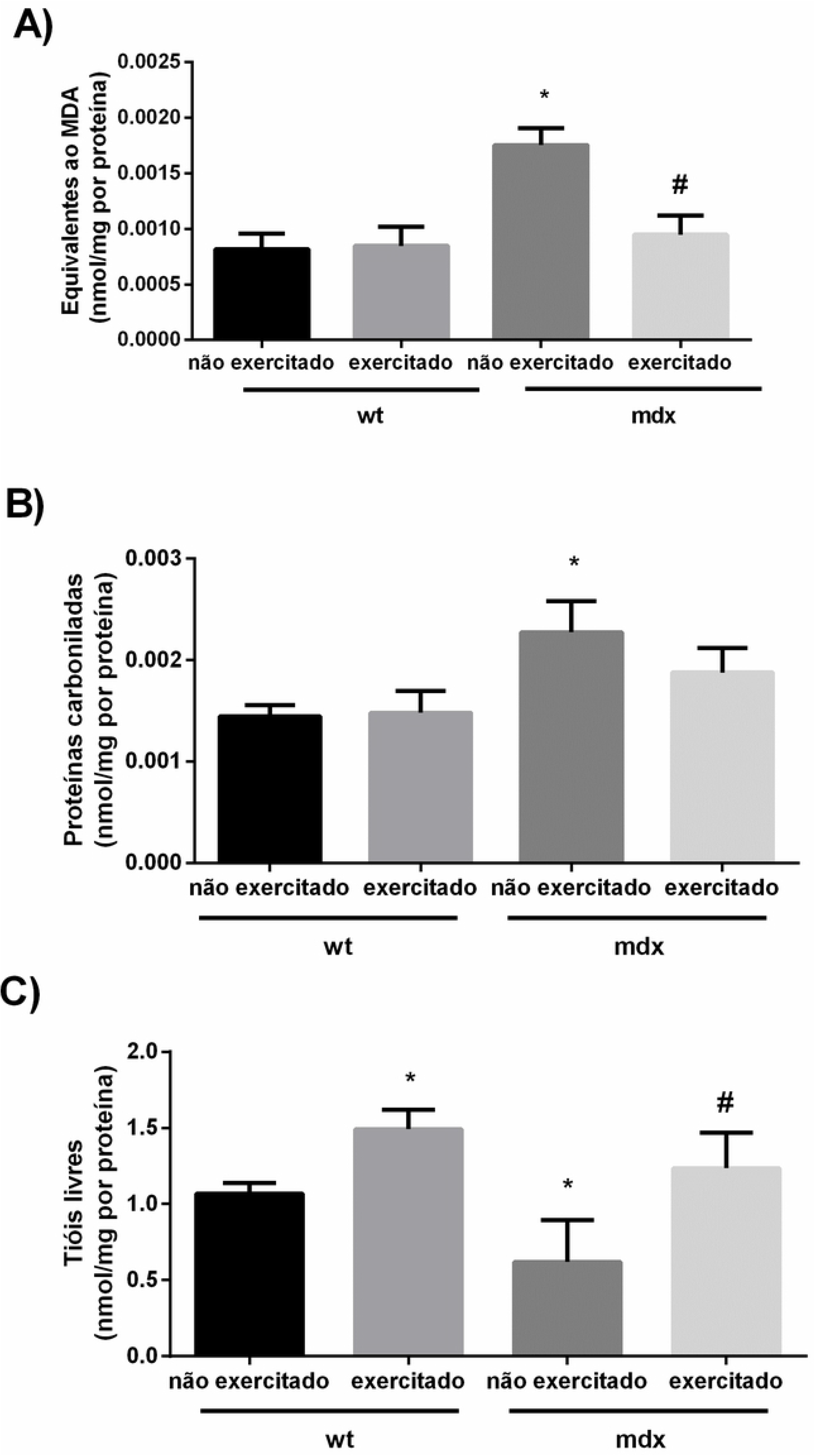
Effect of a swimming protocol on oxidative stress on mouse diaphragm muscle. (* when compared to the non-exercised wt group; ** when compared to the non-exercised mdx group).

The non-exercised mdx animals showed a significant increase in diaphragm lipid peroxidation when compared to the non-exercised wild animals group (*p* <0.05). After the experimental protocol, the mdx animals showed significantly lower lipid peroxidation levels when compared to the non-exercised mdx animals (*p* <0.05), demonstrating that the four-week swimming protocol was able to prevent increased peroxidation. observed in the diaphragm muscle of mdx animals (Figure 6A). Figure 6B shows the result of the evaluation of protein carbonylation in diaphragm. The non-exercised mdx animals showed significantly higher levels of protein carbonation in diaphragm when compared to the non-exercised wild animals (*p* <0.05). However, the mdx animals that underwent the experimental protocol did not show a significant reduction of these levels when compared to the non-exercised mdx animals, showing that the experimental protocol used in this study did not protect against the increase of diaphragm protein carbonylation in mdx animals. The quantitation of free diaphragm thiols is shown in Figure 6C. It can be observed that there was a significant increase in the amount of diaphragm free thiols in the group of wild animals submitted to the experimental protocol, when compared to the non-exercised wild animals (*p* <0.05). There was also a significant decrease in non-exercised mdx animals when compared to non-exercised wild animals (*p* <0.05). When the mdx animals underwent the experimental protocol, there was an increase in the amount of free thiols when compared to the non-exercised mdx animals, demonstrating that swimming was able to protect this change.

### Assessment of oxidative stress in central nervous system structures

Figure 7 shows the results of using a swimming protocol on lipid peroxidation in the prefrontal cortex (Figure 7A), hippocampus (Figure 7B), striated (Figure 7C), and cortex (Figure 7D).

**Figure 7.**
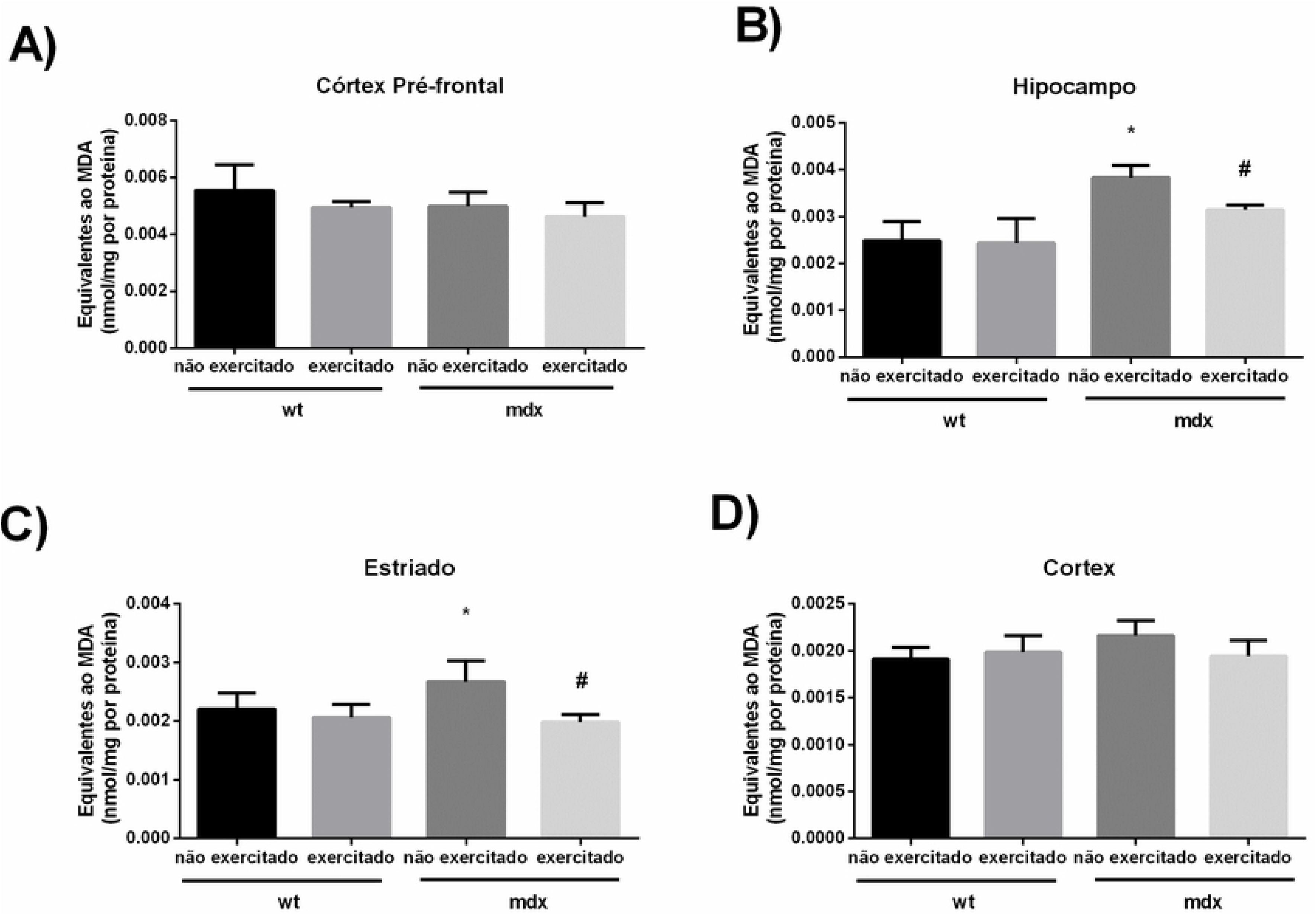
Effect of a swimming protocol on lipid peroxidation in mouse CNS structures. (* when compared to the non-exercised wt group; ** when compared to the non-exercised mdx group).

The non-exercised mdx animals showed a significant increase in hippocampal and striatum lipid peroxidation when compared to the non-exercised wild animals group (*p* <0.05). After the experimental protocol, the mdx animals presented significantly lower lipid peroxidation levels in the hippocampus and striatum when compared to the untrained mdx animals (*p* <0.05), demonstrating that the four-week swimming protocol was able to prevent the increase in lipid peroxidation observed in the hippocampus and striatum structures of mdx animals (Figures 7B and 7C). Figures 7A and 7D show that in the prefrontal cortex and cortex structures there were no significant changes between the analyzed groups. Figure 8 shows the results of using a swimming protocol on protein carbonylation in the prefrontal cortex (Figure 8A), hippocampus (Figure 8B), striated (Figure 8C) and cortex (Figure 8D).

**Figure 8.**
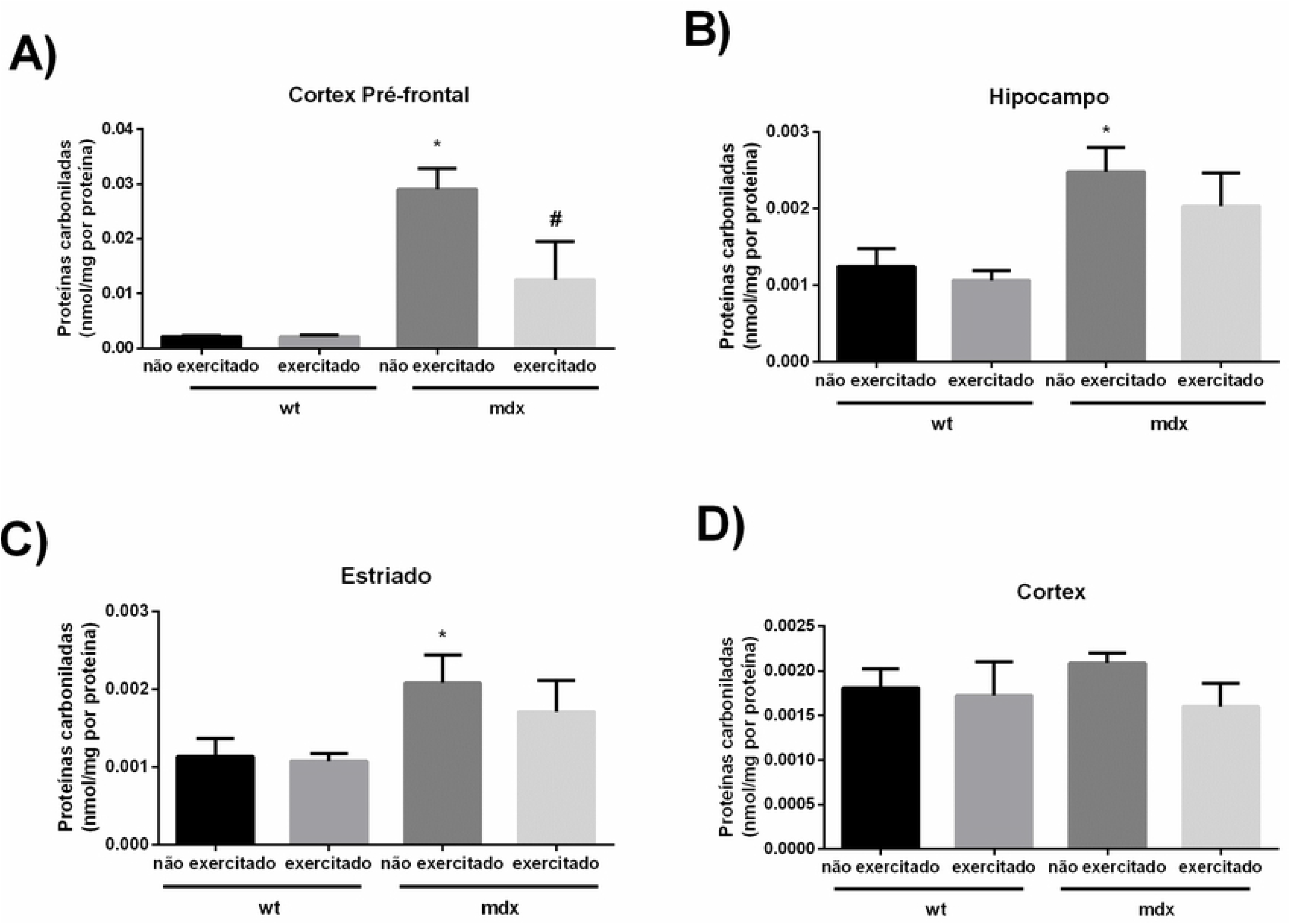
Effect of a swimming protocol on protein carbonylation in mouse CNS structures. (* when compared to the non-exercised wt group; ** when compared to the non-exercised mdx group).

It can be observed that the non-exercised mdx animals showed a significant increase in protein carbonylation in prefrontal, hippocampal and striated cortex when compared to the non-exercised wild animals group (*p* <0.05). After the experimental protocol, the mdx animals showed significantly lower protein carbonylation levels in the prefrontal cortex when compared to the untrained mdx animals (*p* <0.05), demonstrating that the four-week swimming protocol was able to prevent increased protein carbonylation, observed only in the prefrontal cortex structure of mdx animals (Figure 8A). Figures 8B and 8C show that the four-week swimming protocol was unable to prevent increased protein carbonation in the hippocampus and striatum. Figure 8D shows that in the cortex structure there was no significant change between the analyzed groups.

Figure 9 shows the results of using a swimming protocol on free thiols in the prefrontal cortex (Figure 9A), hippocampus (Figure 9B), striated (Figure 9C) and cortex (Figure 9D).

**Figure 9.**
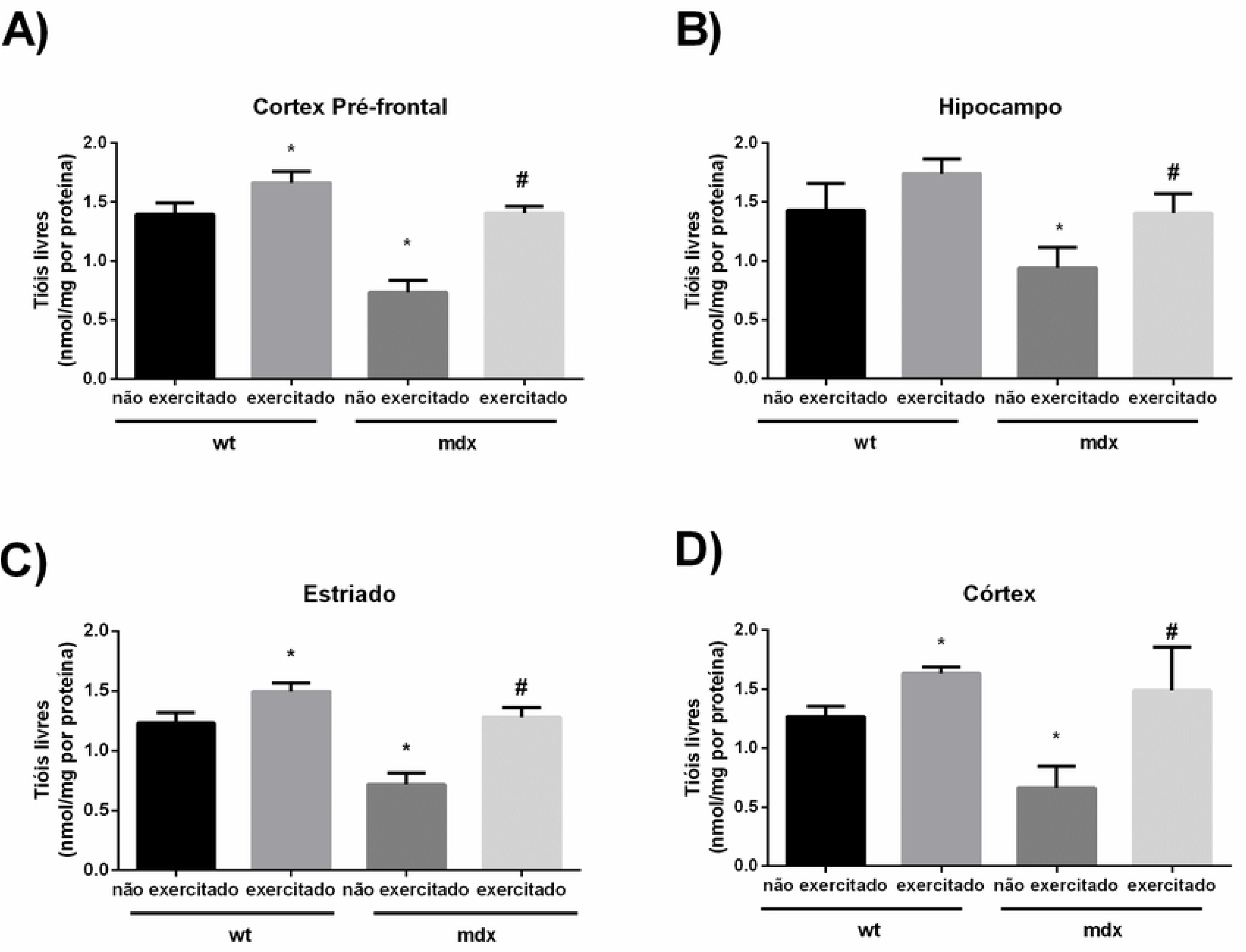
Effect of a swimming protocol on free thiols in mouse CNS structures. (* when compared to the non-exercised wt group; ** when compared to the non-exercised mdx group).

It can be observed that the non-exercised mdx animals showed a significant decrease in free thiols in prefrontal cortex, hippocampus, striatum and cortex when compared to the group of non-exercised wild animals (*p* <0.05). After the experimental protocol, the mdx animals showed significantly higher free thiol levels in prefrontal cortex, hippocampus, striatum and cortex when compared to untrained mdx animals (*p* <0.05), demonstrating that the swimming protocol, by four weeks, it was able to increase antioxidant defenses in all CNS structures of mdx animals. (Figures 9A, 9B, 9C and 9D).

## Discussion

This study aimed to evaluate the effects of swimming on memory and oxidative stress in skeletal muscle and brain tissue in an animal model of Duchenne muscular dystrophy. For this, we used a moderate intensity swimming protocol performed four times a week for four consecutive weeks. The results showed that swimming prevented the aversive memory and habituation impairment in mdx mice. Parallel to this effect, an increase of protein carbonylation in the prefrontal cortex, hippocampus, striatum, diaphragm, and gastrocnemius, and increased lipid peroxidation in the hippocampus, striatum, diaphragm, and gastrocnemius were also observed by the analysis of the parameters related to the oxidative damage, concomitant with the decrease in free thiols in non-exercised mdx animals, evidencing oxidative stress. Interestingly, low intensity swimming was able to prevent oxidative stress in the gastrocnemius and hippocampal and striated structures of these animals. This same protocol increased free thiols in the gastrocnemius, diaphragm, and CNS structures analyzed.

The exercise of swimming (aerobic exercise) used in this study was of moderate intensity, according to the lactate measurements made in the animals. For this, one ml of shampoo was used in the whole container adapted for swimming. Regarding exercise intensity, in addition to the classification based on VO2max and maximum heart rate measurement, which classifies exercise as mild/ low intensity, moderate intensity, or high intensity [17], there is also the model proposed by Gaesser and Poole (1996) proposing three domains relative to the intensity of effort: moderate; heavy; and severe. The moderate domain comprises all the exertion intensities that can be performed without blood lactate modification in relation to the resting values, that is, below the lactate threshold (LL). The heavy domain starts from the lowest effort intensity where lactate rises, and has as its upper limit an intensity corresponding to an average of 4 mM lactate. In the severe domain, there is no stable phase of blood lactate, with blood lactate rising throughout the exercise time, until the point of individual exhaustion [18].

DMD is characterized by an absence of dystrophin protein in skeletal muscle5. However, the literature also shows that dystrophin is absent in brain tissue, and this change is associated with other changes such as oxidative stress [19]. The absence of dystrophin and oxidative stress in the CNS makes cognitive impairment part of the pathophysiology of this disease. In this study, it was evidenced that the swimming protocol was able to protect the aversive memory and habituation memory impairment in the mdx mice submitted to the protocol.

The protocol of the present study was started with 28-day-old animals and ended with animals at 56 days old. At this age, these animals would be expected to have impaired memory and learning, which does not occur in animals submitted to swimming. Thus, it can be said that exercise prevents memory and learning deficits. Although there are no studies relating swimming directly to the prevention of memory impairment and DMD-related learning, there are reports of benefits of swimming practice in cognitive aging [20]. Another study, still with middle-aged animals, showed the benefits of swimming in object recognition memory tests, when combined with dietary supplementation, demonstrating improvement in short and long-term memory [21].

Besides cognitive impairment, another objective of this study was to verify if swimming can alter oxidative stress in neuronal and skeletal muscle tissue. Studies show that oxidative stress is present in the pathophysiological process of DMD, as there is an imbalance between the formation of oxidizing agents and antioxidant activity [22–24]. Oxidative stress is present in DMD, skeletal muscle, and also in the CNS [22]. One of the effects of exercise is increased antioxidant activity. Exercise produces ROS, which act as signals of molecular events, which regulate adaptations in muscle cells, such as the regulation of antioxidant enzymes [25]. In this study, it was observed that swimming increased free thiols, demonstrating that physical exercise is able to increase antioxidant glutathione levels, in agreement with previous studies that demonstrated the protective role of exercise [26,27]. Free thiols are an indirect measure of glutathione activity, an antioxidant present in greater numbers in the CNS [28].

In addition to verifying the influence of swimming on brain impairment, the evaluation of some skeletal muscle tissues is necessary. Since DMD is an essentially neuromuscular disease, it is necessary to include skeletal muscle assessments (gastrocnemius and diaphragm), since the main characteristic of this disease is related to the involvement of these structures with calf pseudohypertrophy, falls, frequent gait loss, and cardiorespiratory dysfunction [29]. Evidence suggests that oxidative stress is associated with aggravation of both respiratory and muscular pathology in these patients [22]. In addition, there are studies showing that exercise can protect against oxidative stress in skeletal muscle in mdx [30,12] mice. The protocol used in this study protected against increased lipid peroxidation in the gastrocnemius in mdx animals. Swimming for four weeks was able to prevent the increased protein carbonylation observed in the gastrocnemius muscle of mdx animals. This finding is in agreement with a study conducted in 2015 which used a swimming protocol and found a decrease in protein carbonylation [12].

The same occurred with the diaphragm muscle, in which the mdx animals had significantly lower lipid peroxidation levels when compared to the non-exercised mdx animals, demonstrating that the four-week exercise protocol was able to prevent increased lipid peroxidation. However, swimming was not able to reverse the increased protein carbonation in the diaphragm. Diaphragm degeneration is a major contributor to dystrophic right ventricular pathology. One study has concluded that although swimming is beneficial for skeletal muscle and increases cardiac function, a 60-min protocol, six days a week, for two months, exacerbates diaphragm degeneration and increases dystrophic phenotype, and causes pulmonary hypertension [30].

Considering the above, it can be suggested, having seen the increase in free thiol formation shown in the results of this study, that the moderate intensity aerobic exercise of swimming may reduce oxidative stress by increasing antioxidant defenses, such as glutathione. However, as there is no consensus in the literature on exercise volume, frequency, and intensity in the treatment of DMD, this study may help to propose new perspectives on the therapeutic use of exercise in the treatment of this disease.

## Conclusion

A swimming protocol applied in mdx mice was able to prevent memory damage and oxidative stress in gastrocnemius and in most of analysed SNC structures with a significant increase in antioxidant activity in all analysed structures.

## Acknowledgments

To research group for their commitment and dedication.

## References

1. Matthew PW. The muscular dystrophies. Continuum Journ 2013;19(6):1535–70.

2. Ljubicic V, Burt M, Jasmin BJ. The therapeutic potential of skeletal muscle plasticity in Duchenne muscular dystrophy: phenotypic modifiers as pharmacologic targets. The FASEB Journ 2014;28:548–68.

3. Savino W, Pinto-Mariz F, Mouly V. Flow Cytometry-Defined CD49d Expression in Circulating T-Lymphocytes Is a Biomarker for Disease Progression in Duchenne Muscular Dystrophy. In Duchenne Muscular Dystrophy. New York, NY: Humana Press; 2018. p. 219–27.

4. Gevaerd MS, Domenech SC, Junior NGB, Higa DF, Lima-Silva AE. Alterações fisiológicas e metabólicas em indivíduo com distrofia muscular de Duchenne durante tratamento fisioterapêutico: um estudo de caso. Fisiot Mov 2010;23(1):93–103.

5. Le Rumeur E. Dystrophin and the two related genetic diseases, Duchenne and Becker muscular dystrophies. Bosn J Basic Med Sci 2015;15(3):14–20.

6. Alemdaroğlu Đ, Karaduman A, Yilmaz ÖT, Topaloğlu H. Different types of upper extremity exercise training in duchenne muscular dystrophy: effects on functional performance, strength, endurance, and ambulation. Muscle & Nerve 2015;51(5):697–705.

7. Hyzewicz J, Ruegg UT, Takeda S. Comparison of experimental protocols of physical exercise for mdx mice and duchenne muscular dystrophy patients. J Neuromuscul Dis 2015;2:325–42.

8. Jansen M, Alfen NV, Geurts ACH, Groot IJM. Assisted bicycle training delays functional deterioration in boys with duchenne muscular dystrophy: the randomized controlled trial “No Use Is Disuse”. Neurorehabilit and Neural Repair 2013;27(9):816–27.

9. Mankodi A, Azzabou N, Bulea T, Reyngoudt H, Shimellis H, Ren Y, et al. Concentric exercise effects on skeletal muscle water T2 in Duchenne muscular dystrophy (P2. 131). Neurology 2017; 88(16 Supplement):P2–131.

10. Nikolai, A. L., Novotny, B. A., Bohnen, C. L., Schleis, K. M. & Dalleck, L. C. Cardiovascular and Metabolic Responses to Water Aerobics Exercise in Middle-Aged and Older Adults. J of Phys Act & Health. 2009; 6,(3): 333–338.

11. Chase, N. L., Sui, X. & Blair, S. N. Comparison of the Health Aspects of Swimming With Other Types of Physical Activity and Sedentary Lifestyle Habits. Int. J. of Aquatic Res & Educ. 2008; 2, (2): 151–161.

12. Hyzewicz, J. Tanihata J, Kuraoka M, Ito N, Miyagoe-Suzuki Y, Takeda S. Low intensity training of mdx mice reduces carbonylation and increases expression levels of proteins involved in energy metabolism and muscle contraction. Free Radic. Biol. Med. (2015); 82, 122–36.

13. Mazzardo-Martins, L. Martins DF, Marcon R, Dos Santos UD, Speckhann B, Gadotti VM, Sigwalt AR, Guglielmo LG, Santos AR. High-intensity extended swimming exercise reduces pain-related behavior in mice: involvement of endogenous opioids and the serotonergic system. J. Pain. 2010; 11, >1384–93.

14. Ishii H, N. Y. Effect of lactate accumulation during exercise-induced muscle fatigue on the sensorimotor cortex. J Phys Ther Sci. 2013; 25, 1637–42.

15. Leussis, M. P. & Bolivar, V. J. Habituation in rodents: A review of behavior, neurobiology, and genetics. Neurosc and Biobehav Rev. 2006; 30, 1045–1064.

16. Ogihara, C. A. Schoorlemmer GH, Lazari Mde F, Giannocco G, Lopes OU, Colombari E, Sato MA. Swimming Exercise Changes Hemodynamic Responses Evoked by Blockade of Excitatory Amino Receptors in the Rostral Ventrolateral Medulla in Spontaneously Hypertensive Rats. BioMed Research International. 2014; 2014.

17. Schubert MM, Washburn RA, Honas JJ, Lee J, D. J. Exercise volume and aerobic fitness in young adults: the Midwest Exercise Trial-2. Springerplus. 2016; 5, (1): 183.

18. Gaesser GA, P. D. The slow component of oxygen uptake kinetics in humans. Exerc Sport Sci Rev. 1996; 24, 35–71.

19. Comim, C. M. Tuon L, Stertz L, Vainzof M, Kapczinski F, Quevedo J. Striatum brain-derived neurotrophic factor levels are decreased in dystrophin-deficient mice. Neuroscience Letters. 2009; 459, (2) 66–68.

20. De Araujo, G. G. Papoti M, Dos Reis IG, de Mello MA, Gobatto CA. Physiological responses during linear periodized training in rats. Eur. J. Appl. Physiol. 2012; 112, (13): 839–852.

21. Cechella, J. L., Leite, M. R., Gai, R. M. & Zeni, G. The impact of a diphenyl diselenide-supplemented diet and aerobic exercise on memory of middle-aged rats. Physiol Behav. 2014; 135, 125–129.

22. Lawler, J. M. Exacerbation of pathology by oxidative stress in respiratory and locomotor muscles with Duchenne muscular dystrophy. J Physiol. 2011; 589, (9): 2161–2170.

23. Lh, M. Bollineli RC, Mizobuti DS, Silveira Ldos R, Marques MJ, Minatel E. Effect of N-acetylcysteine plus deferoxamine on oxidative stress and inflammation in dystrophic muscle cells. Redox Report. 2015; 20, (3): 109115.

24. Kozakowska, M., Pietraszek-Gremplewicz, K., Jozkowicz, A. & Dulak, J. The role of oxidative stress in skeletal muscle injury and regeneration: focus on antioxidant enzymes. J. Muscle Res. Cell Motil. 2015; 36, 377–393.

25. Gomez-Cabrera, M. C., Domenech, E. & Viña, J. Moderate exercise is an antioxidant: Upregulation of antioxidant genes by training. Free Radic. Biol. Med. 2008; 44, 126–131.

26. Antoncic-Svetina, M. et al. Ergometry induces systemic oxidative stress in healthy human subjects. Tohoku University Medical Library. 2010; 221, 43–48.

27. 11. Jia, B. et al. The effects of long term aerobic exercise. Clin. Hemorheol. Microcirc. 2012; 51, 117–127.

28. Halliwell, B. Reactive oxygen species and the central nervous system. J Neurochem. 1992; 59, 609–23.

29. Okama, L. O. et al. Functional and postural evaluation in duchenne and becker muscular dystrophies. Cons e Saúde. 2010; 9,(4) 649–658.

30. Barbin, I. C. C. Pereira JA, Bersan Rovere M, de Oliveira Moreira D, Marques MJ, Santo Neto H. Diaphragm degeneration and cardiac structure in mdx mouse:Potential clinical implications for Duchenne muscular dystrophy. J. Anat. 2016; 228 (5): 784–791.

